# *C. difficile* may be overdiagnosed in adults and is a prevalent commensal in infants

**DOI:** 10.1101/2022.02.16.480740

**Authors:** Pamela Ferretti, Jakob Wirbel, Oleksandr M Maistrenko, Thea Van Rossum, Renato Alves, Anthony Fullam, Wasiu Akanni, Christian Schudoma, Anna Schwarz, Roman Thielemann, Leonie Thomas, Stefanie Kandels, Rajna Hercog, Anja Telzerow, Ivica Letunic, Michael Kuhn, Georg Zeller, Thomas SB Schmidt, Peer Bork

## Abstract

*Clostridioides difficile* is an urgent threat in hospital-acquired infections world-wide, yet the microbial composition associated with *C. difficile*, in particular in *C. difficile* infection (CDI) cases, remains poorly characterised. To investigate the gut microbiome composition in CDI patients, we analysed 534 metagenomes from 10 publicly available CDI study populations. We then tracked *C. difficile* on a global scale, screening 42,900 metagenomes from 253 public studies. Among the CDI cohorts, we detected *C. difficile* in only 30% of the stool samples from CDI patients. However, we found that multiple other toxigenic species capable of inducing CDI-like symptomatology were prevalent. In addition, the majority of the investigated studies did not adhere to the recommended guidelines for a correct CDI diagnosis.

In the global survey, we found that *C. difficile* prevalence, abundance and biotic context were age-dependent. *C. difficile* is a rare taxon associated with reduced diversity in healthy adults, but common and associated with increased diversity in infants. We identified a group of species co-occurring with *C. difficile* exclusively in healthy infants, enriched in obligate anaerobes and in species typical of the healthy adult gut microbiome. *C. difficile* in healthy infants was therefore associated with multiple indicators of healthy gut microbiome maturation.

Our analysis raises concerns about potential CDI overdiagnosis and suggests that *C. difficile* is an important commensal in infants and that its asymptomatic carriage in adults depends on microbial context.

## Introduction

*Clostridioides difficile (*previously *Clostridium* or *Peptoclostridium difficile)* was first isolated in 1935 from the stools of healthy infants (Hall and O’toole 1935), but later identified as a causative agent of pseudomembranous colitis (Bartlett et al. 1978), a severe condition with potentially life-threatening outcomes. While not all *C. difficile* strains are toxigenic, *C. difficile* toxin A (TcdA) and toxin B (TcdB) can increase intestinal permeability and promote intense inflammation (Voth and Ballard 2005), a condition referred to (Bartlett et al. 1978)as *C. difficile* infection (CDI). CDI is typically characterised by diarrhoea and/or pseudomembranous colitis and often unresponsive to repeated antibiotic treatments. Known risk factors include hospitalisation, age (≥65 years of age), and broad-spectrum antibiotic therapy (Eze et al. 2017). CDI has an increasing clinical relevance (at an estimated annual economic burden of US$6.3 billion in the US alone (Bartlett et al. 1978)) and is considered one of the most urgent threats in hospital-acquired infections(CDC, 2019).

*C. difficile* has primarily been studied as a pathogen, with a strong focus on CDI aetiology (Smits et al. 2016), mechanisms of *C. difficile* toxicity (Voth and Ballard 2005) or efficacious therapies (Kociolek and Gerding 2016). However, an increasing number of other enteropathogens has been linked to antibiotics-associated diarrhoea (AAD) with similar, and sometimes indistinguishable, symptomatology to CDI (Tang et al. 2016; Chia et al. 2018; Larcombe et al. 2018), making the correct diagnosis, estimation and management of CDI particularly challenging.

Far less is known about asymptomatic *C. difficile* carriage among the healthy population, as sentinel studies are often limited in sample size and geography (Galdys et al. 2014; Kato et al. 2001; Cui et al. 2021; Mani et al. 2023) (Galdys et al. 2014; Kato et al. 2001). Gut microbial composition associated with *C. difficile* presence in healthy subjects, in particular in infants where *C. difficile* is highly prevalent (Rousseau et al. 2012), remains poorly understood.

Here we leverage public shotgun metagenomic data to characterise gut microbial signatures associated with CDI in adult and elderly patients and to quantify other AAD-associated species with similar signatures. We then contextualise our (Rousseau et al. 2012) analysis by tracing *C. difficile* in a broad multi-habitat collection of 42,900 metagenomes. We investigate *C. difficile* prevalence, relative abundance and its associated microbial community structure and composition in the human gut over lifetime, with particular focus on healthy infants, where several lines of evidence point to *C. difficile* being a commensal and a hallmark species of healthy infant gut microbiome maturation.

## Results

### *C. difficile* is metagenomically detectable in only 30% of patients with a CDI diagnosis

We conducted a metagenomic meta-analysis of the gut microbiome associated with CDI, analysing 534 fecal samples across 10 geographically diverse study populations (Vincent et al. 2016; Smillie et al. 2018; Podlesny and Florian Fricke; Kim et al. 2020; Langdon et al. 2021; Monaghan et al. 2020; Stewart et al. 2019; Kumar et al. 2017; Watson et al.), including patients diagnosed with CDI, diseased control subjects without CDI (D-Ctr, see Methods), and healthy subjects (H-Ctr) (**Figure 1A** and **Supplementary Table 1**).

**Figure 1.**
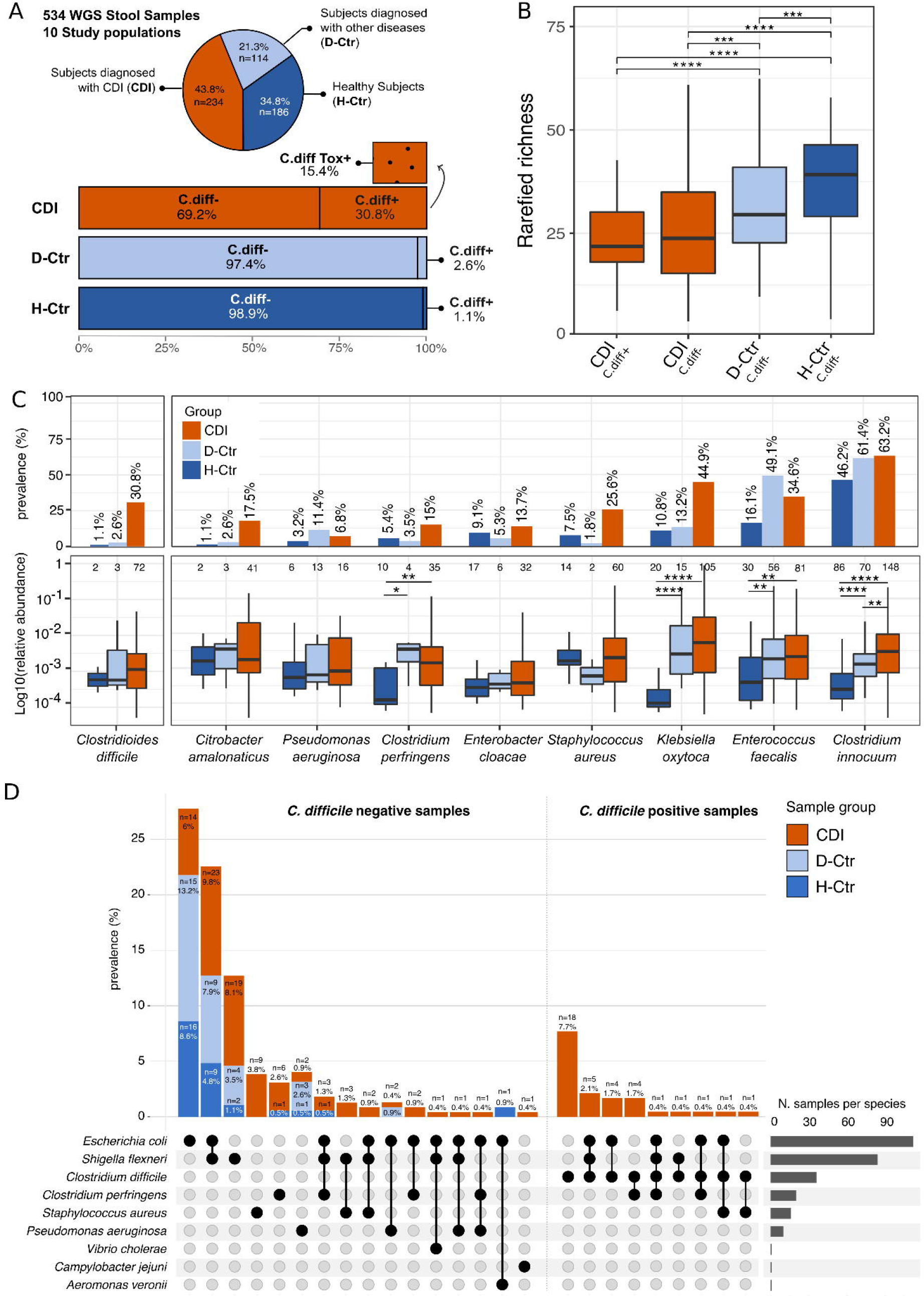
Metagenomic meta-analysis of the gut microbiome associated with CDI. (A) On top, fecal metagenomes used in the meta-analysis from 10 public CDI or diarrhoeal study populations, divided by groups: subjects diagnosed with CDI (CDI, n=234), subjects diagnosed with other diseases than CDI (D-Ctr, n=114), and healthy subjects (H-Ctr, n=186) samples. See Methods for further group description. At the bottom, fraction of *C. difficile* positive samples per group. *C. difficile* toxin genes were found only among CDI *C. difficile* positive samples. (B) Rarefied species richness across the three groups. (C) Prevalence, relative abundance of species that induce antibiotic-associated diarrhoea (AAD) among CDI and controls. (D) Prevalence of single and multiple toxigenic species found among *C. difficile* positive and *C. difficile* negative samples, divided by sample group. Mean comparison p-values calculated using Wilcoxon-test, * for P ≤ 0.05, ** for P ≤ 0.01, *** for P ≤ 0.001, **** for P ≤ 0.0001, empty when non significant.

Using mOTUs-based species-level taxonomic profiling, *C. difficile* was detectable in just 30% of CDI samples at established metagenomic detection limits (Milanese et al. 2019) (**Figure 1A**), with considerable variance across study populations (9.2%-92.3%), and with much lower prevalence among diseased (2.6%) and healthy (1.1%) controls (**Supplementary Figure 1A**). *C. difficile* toxin genes *tcdA* and *tcdB* were detectable in about half of *C. difficile*-positive CDI metagenomes, but not in any *C. difficile*-negative CDI samples nor controls, indicating a lower sensitivity of toxin-based detection (**Figure 1A**, see Methods). We confirmed that marker gene-based *C. difficile* detection was not limited by sequencing depth (linear mixed-effect model p=0.35, ANOVA, adjusted p=0.54; R^2^=0.01), and that detection was not skewed towards metagenomic detection limits (Parks et al. 2021). Neither species richness (**Figure 1B**) nor composition (**Supplementary Figure 2**) differed between *C. difficile*-positive and -negative CDI samples, suggesting that *C. difficile* presence was not associated with characteristic community-level shifts in these generally disbalanced microbiomes.

### The microbiome of CDI patients is characterised by an enrichment of enteropathogens beyond *C. difficile*

Several other species were enriched in prevalence or abundance in CDI patients relative to controls (**Figure 1C, Supplementary Figure 3**). Among these, we identified several opportunistic enteropathogens known to induce antibiotic-associated diarrhoea (AAD), resulting in a CDI-like diarrhoeal symptomatology (Chia et al. 2018; Chia et al. 2017; Kiu and Hall 2018; Zollner-Schwetz et al. 2008; Larcombe et al. 2016; Larcombe et al. 2018; Högenauer et al. 1998; Fakhkhari et al. 2022), such as *Staphylococcus aureus, Klebsiella oxytoca, Enterococcus faecalis, Enterobacter cloacae, Citrobacter amalonaticus, Clostridium innocuum, Clostridium perfringens*, and *Pseudomonas aeruginosa*. In particular, *C. innocuum* was highly prevalent (63.2%) in CDI samples, and significantly enriched relative to both healthy and diseased controls (**Figure 1C**, Wilcoxon test p=8.2×10^-16^ and p=4.3×10^-3^, respectively). 94% of all CDI samples contained at least one of these other AAD species (*C. difficile* excluded) with variable prevalence across study populations (Fisher’s test, p=2.2×10^-16^, OR=29.62; **Supplementary Figure 1A and 1B**). Presence and potential toxigenicity of several enteropathogens was further confirmed by the detection of characteristic toxin genes that were likewise enriched in CDI samples (**Figure 1D**), in particular enteropathogenic *E. coli* and *S. flexneri* toxin genes (see Methods for full list). Overall, CDI samples, independently of *C. difficile* presence, were significantly enriched (p=2.2×10^-16^) in species carrying toxin genes in terms of cumulative relative abundance, indicating that CDI symptomatology could be entirely or partially driven by other enteropathogens than *C. difficile*.

To integrate these univariate associations of individual species into characteristic and predictive multi-species signatures of CDI, we next trained a series of LASSO-regularised logistic regression models in a leave-one-study-out validation approach (see Methods). As expected, *C. difficile* was the most predictive species for CDI samples, although the enrichment of other putative AAD enteropathogens (such as *S. aureus* or *C. perfringens*) was also highly characteristic, in particular in CDI samples where *C. difficile* was not detectable (**Figure 2A**). Moreover, several oral species (such as *Veillonella parvula, Veillonella atypica* or *Rothia dentocariosa*) were enriched in CDI gut metagenomes, implying increased oral-intestinal microbial transmission(Schmidt et al., 2019) among these patients. Enrichment of common probiotics (*Lactobacillus casei, L. plantarum* or *L. fermentum*) in CDI is likely due to the widespread use of probiotic therapy among these patients (Golić et al. 2017; Na and Kelly 2011).

**Figure 2.**
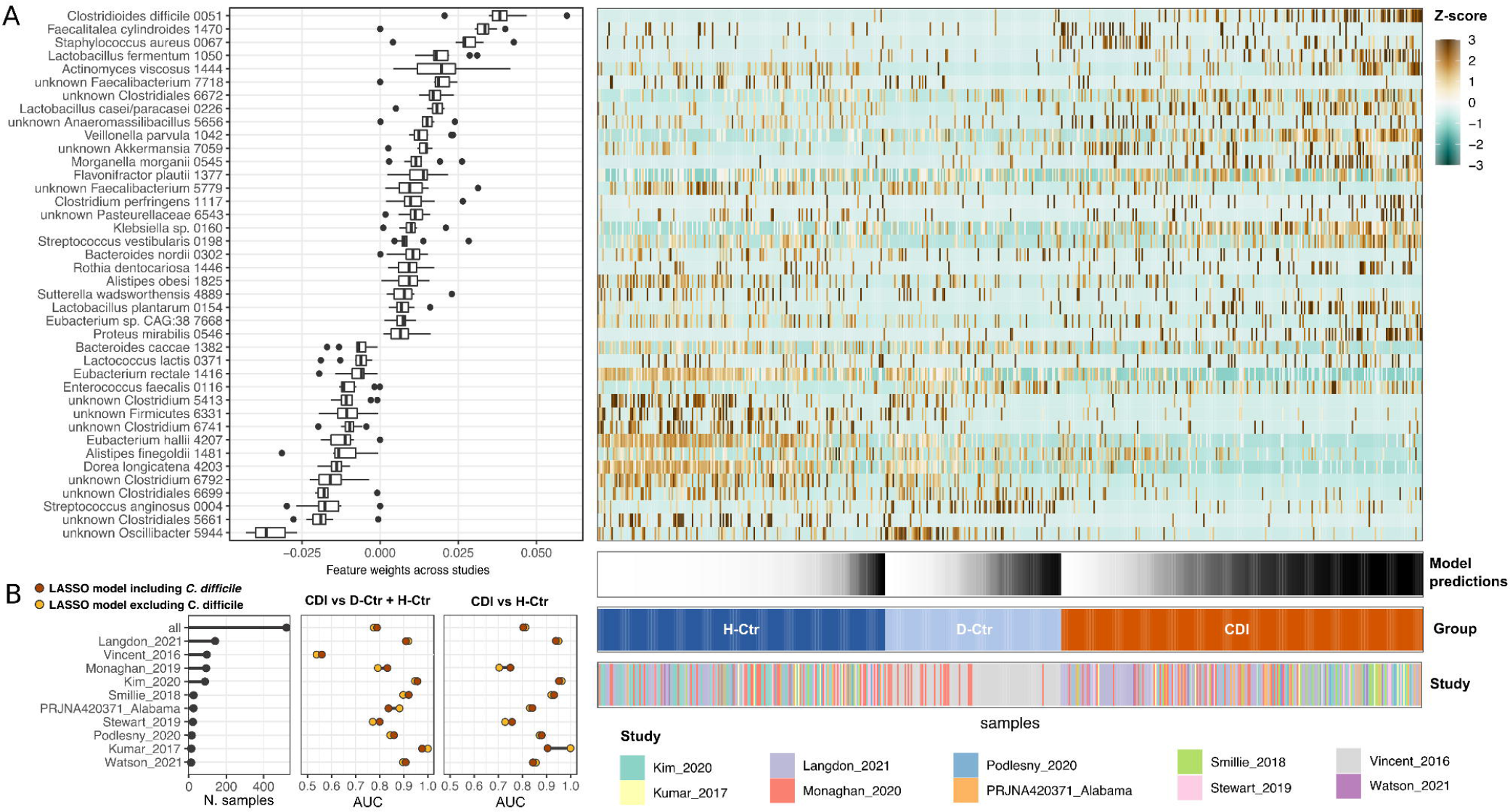
A CDI-specific microbial signature. (A) Microbial species signature associated with CDI, healthy controls and diseased controls, as seen by LASSO model and (B) its associated AUC values for all samples as well as for single study populations, when comparing CDI to healthy and diseased controls combined (left) and CDI with healthy controls only (right). In the latter, the AUC value of Vincent_2016 is left intentionally blank as only diseased controls (D-Ctr) are available for this study population (see **Supplementary Table 1**).

Using these models, CDI samples were distinguishable from diseased and healthy controls with high accuracy, at an area under the receiver operating characteristics curve (AUROC) of 0.78, ranging between 0.56 and 0.98 depending on the study population (**Figure 2B**), in line with a previous 16S-based investigation (Schubert et al. 2014). Model performance moderately improved when considering only healthy controls (overall AUROC = 0.81) (**Figure 2B**). To assess the importance of *C. difficile* to predicting CDI state, we next re-trained models under explicit exclusion of *C. difficile*. Interestingly, this did not lead to a noticeable reduction in accuracy (**Figure 2B**), indicating that the presence or enrichment of *C. difficile* was indeed not an essential defining feature of CDI samples. Rather, *C. difficile* appeared to be just one of several AAD species characterising the CDI-associated microbiome in varying constellations, forming a loose clique instead of fixed co-occurring species groups.

### Extensive asymptomatic carriage of *C. difficile* across age and geography

Given that *C. difficile* appears to have a much lower prevalence in CDI patients than anticipated, we next studied its prevalence in an extended collection of 253 publicly available studies, including a novel cohort of 447 samples from the citizen science project MyMicrobes, for a total of 42,900 shotgun metagenomic samples (**Supplementary Table 2).** This collection included healthy and diseased subjects of all ages, with samples from multiple body sites, hosts, environments and geographic locations (**Figure 3A and 3B**). Statements from this point onwards refer to this extended collection of samples.

**Figure 3.**
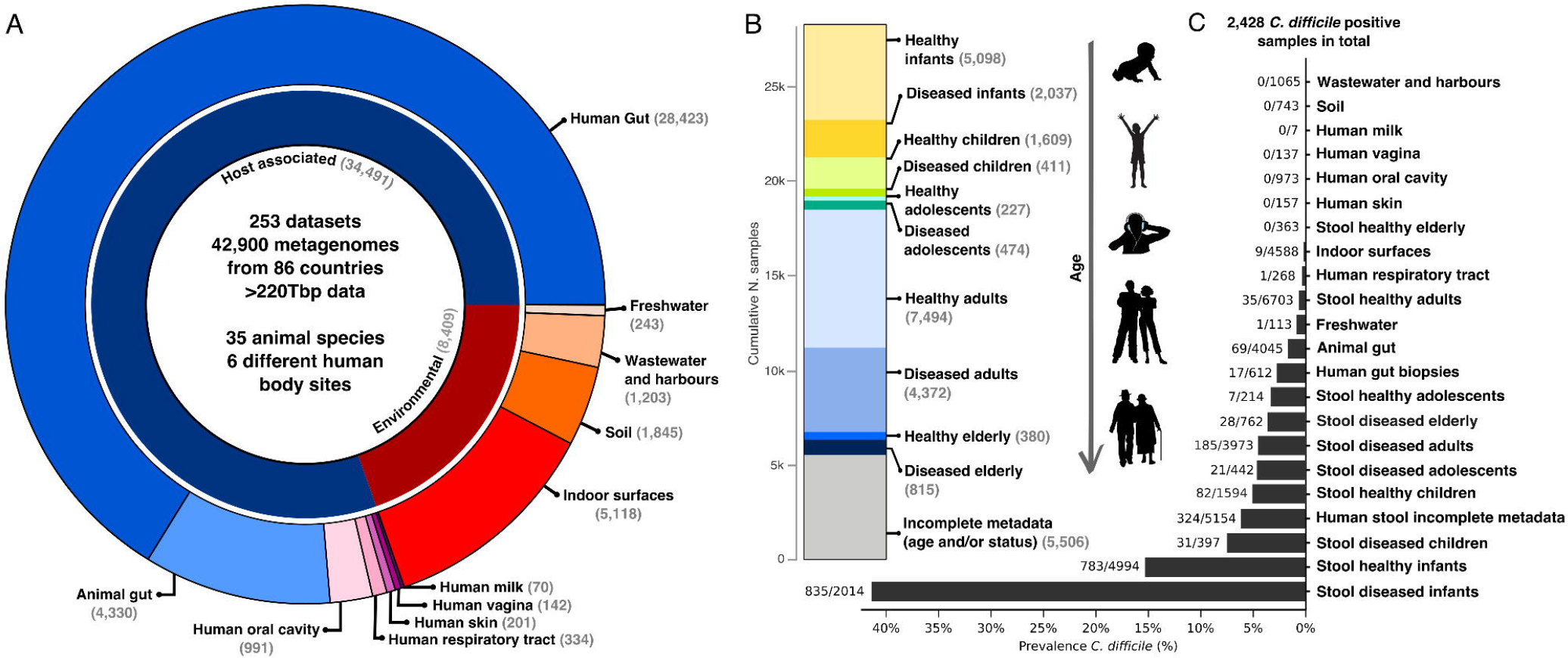
*C. difficile* tracking in a global dataset collection. (A) Overview of the global dataset collection composed of 42,900 samples, from 253 publicly available studies, including host-associated and environmental samples (internal ring), and their subcategories (external ring). The human gut was the most represented environment (66.3%, n=28,423) in our collection. (B) Stratification of the human gut samples, divided by health status and age group (see Methods for detailed age group description, numbers refer to the number of samples prior to read filtering). (C) Prevalence of *C. difficile* positive samples per category. Total values refer to the number of samples after the initial read filtering and before time series dereplication (see Methods). From this point onwards, results refer to this extended collection of 42,900 samples.

*C. difficile* was detected almost exclusively in human and animal feces (8.7% and 1.7% prevalence, respectively) with sporadic detection in hospital surfaces (0.2%), the respiratory tract of diseased patients (0.4%) and freshwater (0.9%) (**Figure 3C** and **Supplementary Table 3**). *C. difficile* was also detectable in the colon, ileum and cecum lumen, and biopsies from the colon and rectal mucosa of healthy subjects. We again confirmed in this larger dataset that there was no relationship between *C. difficile* detection and sequencing depth (linear mixed-effect model p=0.36; logistic regression, ANOVA adjusted p=0.52, **Supplementary Figure 4A and 4B**). Nevertheless, we adjusted all subsequent analyses for total sequencing depth.

In healthy human populations, both *C. difficile* prevalence and relative abundance in fecal samples varied considerably across age ranges. Infants aged one year or younger showed by far the highest *C. difficile* carriage, up to 76.5% prevalence at the age of 8 to 10 months (**Figure 4A**), at an average relative abundance of 1.1% (and up to 50% in a two-weeks-old healthy preterm infant). *C. difficile* prevalence remained elevated (43.4%) between 1 and 4 years of age, in contrast to previous reports (Jangi and Thomas Lamont 2010; Lees et al. 2016). With increasing age, *C. difficile* was increasingly rare, found in less than 1% of the healthy population (6.5% in adolescents, 0.8% in adults) (**Figure 4A** and **Supplementary Table 3**) at average abundances of 0.2%, suggesting a considerably lower asymptomatic carriage among adults than previously estimated (up to 15% (Crobach et al. 2018)). *C. difficile* carriage varied across geography (**Supplementary Figure 5A**) and was lower in non-westernised populations, at 4.8% and 0.4% in infants and adults, respectively.

**Figure 4.**
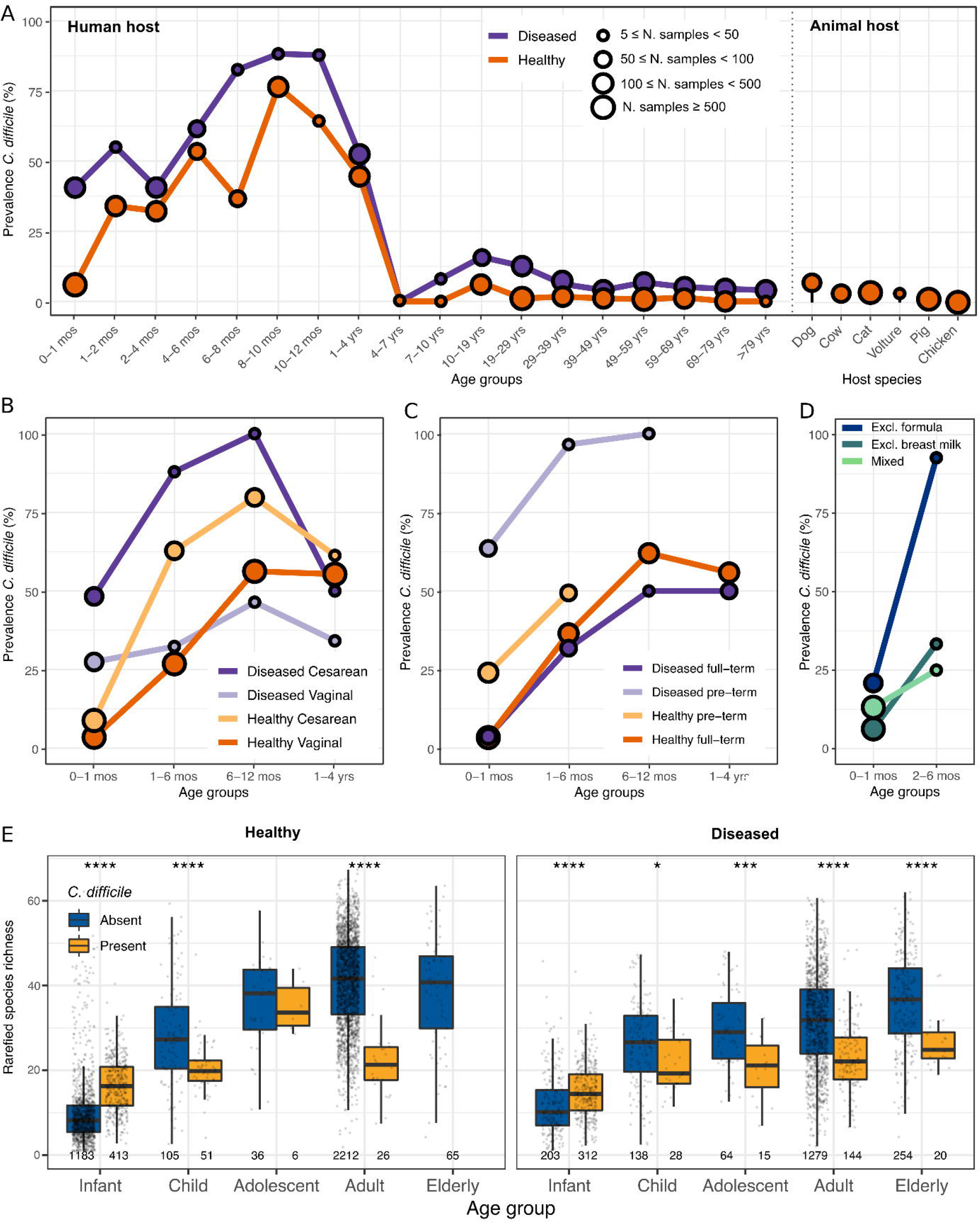
*C. difficile* prevalence and associated microbial diversity. (A) *C. difficile* prevalence in stool samples in healthy and diseased human (left) and animal (right) hosts over lifetime. *C. difficile* prevalence divided by (B) health status and delivery mode, (C) health status and prematurity and (D) feeding mode within the first semester of life. For animal samples, only species with at least ten total samples are shown. (E) Species richness of human stool samples, with and without *C. difficile,* across age groups in healthy (left) and diseased subjects (right). See Methods for details on disease and age group definitions. For samples belonging to time series, only the representative samples are included in the estimations (see Methods). Mean comparison p-values calculated using t-test, * for P ≤ 0.05, ** for P ≤ 0.01, *** for P ≤ 0.001, **** for P ≤ 0.0001, empty when non significant.

### *C. difficile* is enriched upon antibiotic treatment

Diseased or antibiotics-treated subjects exhibited consistently increased *C. difficile* carriage rates across all age groups (**Figure 4A**). Prevalence was highest among infants taking antibiotics (81.1%, relative abundance 2.2×10^-2^) and infants suffering from cystic fibrosis (78.7%, 4.0×10^-3^) or neonatal sepsis/necrotizing enterocolitis (53.7%, 3.2×10^-2^; **Supplementary Figure 5B**). An elevated *C. difficile* asymptomatic carriage rate in cystic fibrosis patients is well documented (Deane et al. 2021) and may be the result of long-term antibiotics usage and frequent exposure to the hospital environment (Bauer et al. 2014). Among adolescents, adults and elderly, *C. difficile* prevalence was highest in patients suffering from CDI (31%, see above) or undefined diarrhoea (25.9%), followed by subjects taking antibiotics (on average 25% across age groups, relative abundance 4.6×10^-3^). Inflammatory bowel diseases, liver diseases, diabetes, and various forms of cancer were likewise associated with an increased *C. difficile* carriage relative to the healthy background population (**Supplementary Figure 5B**).

### *C. difficile* is a frequent commensal in the healthy infant gut

*C. difficile* was among the most prevalent species in infants during the first year of life. Carriage was increased in infants born via C-section (28.7%; logistic regression, adjusted ANOVA p=3.0×10^-3^, logOR=2.0×10^-5^) compared to age- and health status-matched vaginally born infants (19.6%) (**Figure 4B** and **Supplementary Figure 6**). Likewise, prevalence was moderately increased among pre-term (39.14%; p=1.7×10^-1^, logOR=1.1×10^-5^) compared to full-term healthy infants (23.8%) (**Figure 4C** and **Supplementary Figure 6**). Exclusively formula-fed infants were more frequently colonised by *C. difficile* (43.2%; p=5.6×10^-6^, logOR=3.1×10^-5^), compared to partially or exclusively breastfed infants during the first 6 months of life (14.2% and 8.6% respectively) (**Figure 4D**), in line with previous reports (McDonald et al. 2018; Drall et al. 2019).

*C. difficile* was not maternally acquired at birth: across 829 mother-infant pairs included in the dataset, we found no single case of vertical transmission of *C. difficile* from any of the maternal body sites investigated (gut, vagina and oral cavity). Rather, *C. difficile* was likely sourced from the social environment (i.e. other infants, family members or pets) during early life, in line with previous observations on *Clostridia* (Korpela et al. 2018). Leveraging longitudinal data, we observed that the first appearance of *C. difficile* was not arbitrarily distributed over the first year of life, but instead concentrated in two defined time windows, between the 2^nd^-4^th^ and 8^th^-10^th^ months of age (**Supplementary Figure 7**). The first interval coincided with the increased strain influx from environmental sources to the infant gut reported in a previous study (Ferretti et al. 2018), probably due to the start of mouthing. The second interval, dominated by samples from the UK, overlaps with the paid maternity leave length in that country (Thevenon et al. 2016), suggesting that it could be due to the start of day care. C-section and premature birth were associated with increased rate of *C. difficile* appearance, compared to vaginally born and full-term infants (**Supplementary Figure 7**). Together these results suggest that *C. difficile* is acquired from the environment, and that increased exposure to novel environmental sources increases the chances of *C. difficile* acquisition. In addition, during the first year of life, the gut microbiome of healthy infants colonised with *C. difficile* was significantly more similar, in terms of composition, to that of their mothers (**Supplementary Figure 8**).

These vast differences in *C. difficile* carriage across age did not translate into differential toxin burdens. *C. difficile* toxin genes were detectable in metagenomes of healthy *C. difficile*-positive infants (22.4%) and adults (18.6%) at similar rates (**Supplementary Figure 9**).

### *C. difficile* occurs in distinct biotic and physiological contexts in infants and adults

The *C. difficile*-associated microbiome varied considerably across age ranges. *C. difficile* carriage was associated with a significantly decreased overall species richness in subjects above 1 year of age, independently of the health status (**Figure 4E**, see Methods for the definition of diseased). However, in healthy infants we observed the opposite trend: species richness was higher in subjects carrying *C. difficile* (**Figure 4E** and **Supplementary Figure 10A**), irrespective of delivery mode, gestational age or geography (**Supplementary Figure 10B and 10C**). Species richness was also elevated in *C. difficile*-positive diseased infants, albeit to a lesser degree. In line with these trends, *C. difficile* carriage was associated with higher community evenness (lower dominance of individual species) in infants, but a more uneven community structure in adults (**Supplementary Figure 10D**).

To further characterise how the biotic context of *C. difficile* in the gut changes over lifetime, we studied its co-occurrence with other intestinal species, stratified by age group and health status. We broadly distinguished two groups of species based on their co-occurrence relationships with *C. difficile* (**Figure 5**), independently of the presence of its toxin genes. The first set of species (“Group 1”) consistently co-occurred with *C. difficile* across all age groups. Part of this group are *Clostridium paraputrificum* and *Clostridium neonatale* (**Supplementary Figure 11A**), in line with previous reports(Daquigan et al., 2017), an association that was conserved also in animal hosts (**Supplementary Figure 11B**). Group 1 was enriched in facultative anaerobic or aerobic species and comprised several known AAD enteropathogens, such as *K. oxytoca, C. perfringens, C. amalonaticus* or *C. innocuum*. A second set of species (“Group 2”) co-occurred with *C. difficile* exclusively in healthy infants, but showed strong avoidance patterns in later life stages and, to a lesser extent, in diseased infants. Group 2 was characterised by obligate anaerobic species typical of the healthy adult gut microbiome and included several common butyrate producers (such as *Roseburia sp., Faecalibacterium prausnitzii* or *Anaerostipes hadrus*) (**Figure 5** and **Supplementary Figure 12**).

**Figure 5.**
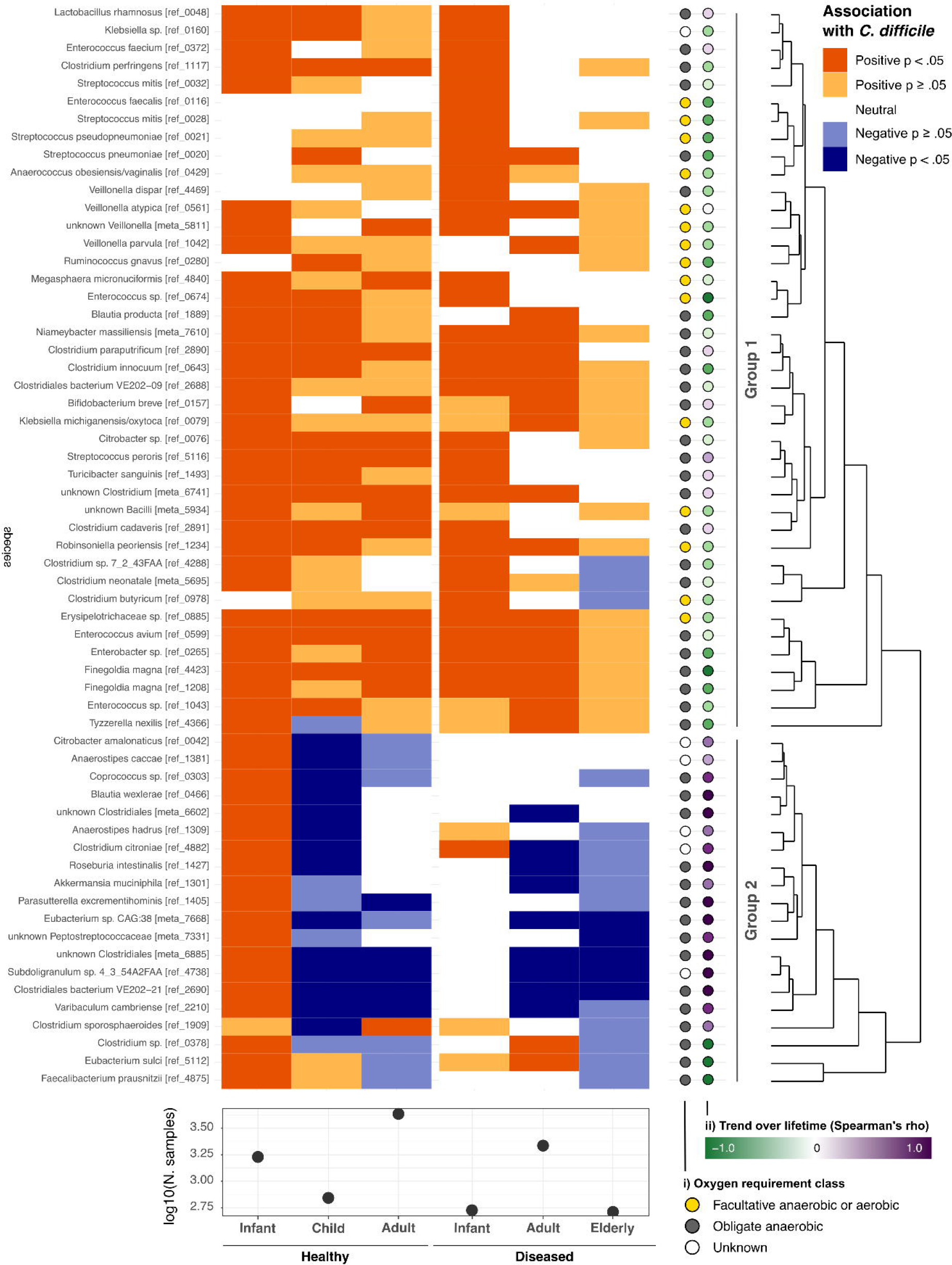
Biotic context associated with *C. difficile*. Species associated with *C. difficile*, divided by age group and health status: significant positive associations are shown in dark orange, significant negative ones in dark blue (p-values adjusted using BH). Lighter shades indicate non significant associations. Only species significantly associated with *C. difficile* in at least one age group are shown. Number of samples per each category shown in the lower part. See Methods for details on age group definitions. Species group separation was not affected by presence of *C. difficile* toxin genes (data not shown). On the right, per species annotation of their i) oxygen requirement (see also Supplementary Figure 12 for the differential enrichment across the two groups) and ii) trend over lifetime. Positive Spearman’s rho values indicate species more commonly found in the gut microbiome of healthy adults, negative values the opposite trend.

To investigate whether healthy infants carried specific *C. difficile* strains, we performed an analysis of metagenomic single nucleotide variants (SNVs). Despite the distinct biotic context associated with *C. difficile* presence in healthy infants at the species level, *C. difficile* strain populations found in this group did not cluster separately from those identified in other age groups and health status, with consistent patterns of toxin-gene carriage (**Supplementary Figure 13**). Therefore, the general asymptomatic nature of *C. difficile* carriage in healthy infants is likely determined by its surrounding microbial community and host age, rather than specific strain groups.

## Discussion

*C. difficile* is predominantly studied in context of its pathogenicity, yet its role in the human gut is arguably much more nuanced. Our data suggests that *C. difficile* may be both overestimated as a pathogen among the elderly and underestimated as a commensal among infants, and that its ambivalent role in the gut is malleable throughout the host’s lifespan depending on external triggers (like antibiotics treatment) and biotic context.

We detected fecal *C. difficile* in just 30% of cases diagnosed with CDI, consistent with previous smaller-scale reports based on 16S rRNA (Daquigan et al. 2017; Seekatz et al. 2016) and WGS metagenomics (Vincent et al. 2016; Zhou et al. 2016), even when combined with laboratory-based tests (Seekatz et al. 2016). This surprisingly low prevalence was neither due to metagenomic detection limits, as sequencing depth was not associated with *C. difficile* prevalence, nor could it be explained by (temporary) reduction of *C. difficile* abundance below detection limits by antibiotics, as stricter inclusion criteria for samples from active and symptomatic CDI cases did not change the observation. A more likely explanation for the under-detection of *C. difficile* among CDI patients is a lack of diagnostic specificity. Challenges in CDI diagnosis are further complicated by the vast array of protocols and diagnostic algorithms available in clinical practice (Jo et al. 2022). Diarrhoeal symptoms, often the initial diagnostic prompt, are not specific to CDI (Bartlett and Gerding 2008; Polage et al. 2012; Jackson et al. 2016; Reich et al. 2019) and pseudomembranous colitis, long thought to be characteristic (and often eponymously referred to as “*C. difficile* colitis”), may indeed be caused by other enteropathogens, such as *C. innocuum* or *E. coli (Gateau et al. 2018; Humphries et al. 2013; Burnham and Carroll 2013; Lee et al. 2021; McDonald et al. 2018; Crobach et al. 2016; Tenover et al. 2011; Rodriguez et al. 2016; Martínez-Meléndez et al. 2017; Litvin et al. 2009)*. Additional challenges include the large variability in sensitivity and specificity in diagnostic laboratory tests (Gateau et al. 2018; Humphries et al. 2013; Burnham and Carroll 2013; Lee et al. 2021; McDonald et al. 2018; Crobach et al. 2016; Tenover et al. 2011; Rodriguez et al. 2016; Martínez-Meléndez et al. 2017; Litvin et al. 2009; BBL™ Clostridium difficile Selective …), contradicting results between cultivation and toxin detection ass(Polage et al. 2015)ays (Parks et al. 2011) and an excessive reliance on single molecular tests (Polage et al. 2015) Recommended guidelines discourage stand-alone tests for CDI diagnosis and recommend a two-step procedure (Crobach et al. 2016; Gateau et al. 2018), but due to costs, test availability, capacity and turnaround times, diagnostic algorithms vary between hospitals and often deviate from recommended procedures (Tenover et al. 2011). Despite the recent improvements in the CDI diagnosis guidelines awareness, low compliance is still reported (Viprey et al. 2023). In our meta-analysis, almost two thirds (62.5%) of studies deviated from recommended best practices for CDI diagnostic protocols.

Moreover, *C. difficile*’s role in causing antibiotics-associated diarrhoea (AAD) or pseudomembranous colitis (PMD) may be overstated, as simultaneous colonisation of multiple enteropathogens is very common among CDI patients (Hensgens et al. 2014; Spina et al. 2015). Enteropathogens like *C. perfingens, C. innocuum, S. aureus, E. coli* or *S. flexneri*, among others, are known to induce AAD or PMC (Chia et al. 2018; Kiu and Hall 2018; Zollner-Schwetz et al. 2008; Larcombe et al. 2018; Högenauer et al. 1998; Larcombe et al. 2016) and were prevalent and enriched among CDI samples, as were their characteristic toxin genes – in particular also in samples where *C. difficile* was not detectable. Indeed, while *C. difficile* was the most individually characteristic species of CDI, its presence was not an essential input to microbiome-based classification models accurately predicting CDI-diagnosed samples. Taken together, this suggests that *C. difficile* is often not the most parsimonious explanation for clinical manifestations of CDI that may instead be driven by alternative enteropathogens, implying mis-or over-diagnosis in practice and an overestimation of the real burden of *C. difficile* relative to other AAD species. Differential diagnosis against multiple enteropathogens may therefore stratify patients with CDI-like symptoms, towards adapted therapeutic interventions.

Widespread asymptomatic carriage in healthy individuals is a further indicator that *C. difficile* plays a complex role in the gut, beyond pathogenicity. We tracked *C. difficile* across an extensive set of 42,900 metagenomes sampled from multiple habitats and hosts, includingn=28,423 (66.3%) human gut samples which covered a wide range of subject ages, disease states and geographic origins. *C. difficile* was particularly common among infants, peaking at >75% prevalence at 8-10 months of age. On average *C. difficile* was present in 25% of healthy infants aged 1 year or younger, in line with previous studies (Lees et al. 2016; Jangi and Thomas Lamont 2010).

In healthy infants, the occurrence of *C. difficile* coincided with an increased resemblance to the maternal gut microbial community, general increase in microbiome richness and with several obligate anaerobic commensals, butyrate producers and a generally more adult-like microbiota, indicating that *C. difficile* may be a transient hallmark of healthy gut microbiome maturation (Bäckhed et al. 2015; Yassour et al. 2016; Ferretti et al. 2018; Chu et al. 2017; Rodríguez et al. 2015). In contrast, *C. difficile* presence in adulthood was associated with decreased microbiome richness and an enrichment in facultative and obligate aerobes, generally associated with dysbiosis (Rigottier-Gois 2013; Rivera-Chávez et al. 2016). We found multiple lines of evidence suggesting that the transition between these apparent dichotomic states (commensal in infancy, pathobiont in adulthood), begins at around 10 (±2) months of age. We hypothesise that *C. difficile* could be an indicator species of a larger-scale and fundamental change taking place in the microbial ecology of the gut.

*C. difficile* toxin genes were likewise prevalent among infants, in line with previous reports (Rousseau et al. 2012; Kubota et al. 2016; Mani et al. 2023), indicating asymptomatic carriage of toxigenic strains. Indeed, *C. difficile* testing is generally discouraged by paediatric guidelines even in presence of diarrhoeal symptoms (McDonald et al. 2018; Schutze et al. 2013) and toxin concentration may not be a reliable proxy for inferring disease and its severity (Davis et al. 2016; Kubota et al. 2016). The apparent protection from CDI symptoms has been ascribed to a lack of toxin receptors in the immature infant gut, although this early hypothesis (Chang et al. 1986) has since been called into question (Keel and Songer 2007; Eglow et al. 1992). The biotic context of *C. difficile* suggests additional or alternative explanations. We observed a significant enrichment in butyrate producers co-occurring with *C. difficile* in healthy infants, but not later in life nor in diseased infants. High levels of butyrate have been linked to inflammation inhibition, regulation of cell-to-cell tight junctions and increased mucin production (and therefore mucosal layer thickness and integrity) (Cornick et al. 2015; Willemsen et al. 2003), all considered protective against *C. difficile* toxins (Pruitt and Lacy 2012).

Our analyses are descriptive and arguably inherently limited by the underlying metagenomic data, such as DNA extraction bias against endospores (Felczykowska et al. 2015) or general detection limits due to finite sequencing depth. Moreover, although some longitudinal data was available, time series were too sparse to infer causal relationships, such as e.g. the possible role of *C. difficile* in a partially deterministic community succession in the infant gut. To our knowledge, ours is the largest single-species metagenomic survey to date, demonstrating the utility of metagenomic meta-studies as a nuanced approach to gut microbial ecology, beyond *C. difficile*. Overall, our results suggest that the ability of *C. difficile* to induce disease may be context-dependent and multifactorial (i.e influenced by the rest of the microbiome composition, toxin presence and host age). Further study of *C. difficile*’s association with health outcomes should therefore adopt a more holistic view of the ambivalent role of this species over the human lifespan.

## Methods

### Human study

We compiled a metagenomic dataset of both public (see below) and 447 newly generated samples. The novel cohort is part of the citizen science-based my.microbes project (http://my.microbes.eu/) (Voigt et al. 2015), conducted in accordance with the WMA Declaration of Helsinki and with the approval of the EMBL Bioethics Internal Advisory Board. Sample collection, processing and sequencing were performed as previously described in (Voigt et al. 2015; Costea et al. 2017; Hildebrand et al. 2021). This cohort includes human fecal (n=437) and oral (n=10) samples, from a total of 142 subjects (aged 0-97 years). Self-reported subject metadata are included in **Supplementary Table 4**.

### Public data overview

#### CDI meta-analysis

In the CDI meta-analysis (see **Supplementary Table 1)**, out of 534 samples in total (from as many subjects, with an average age of 60±19.9 years), 234 were identified as CDI, 114 as diseased controls and 186 as healthy controls. Samples from subjects diagnosed with CDI in the original study population were classified as CDI in our meta-analysis. Samples were identified as controls in any of the following cases: (i) H-Ctr: healthy subject, reported as “control” in the metadata of the original study population; or (ii) D-ctr: diseased subject, that either had diarrhoeal symptoms but negative CDI diagnostic outcome, or was diarrhoea-free but diagnosed for a disease other than CDI. The majority of the samples in the latter group (D-Ctr) belong to hospitalised subjects. Study-specific CDI diagnostic criteria are indicated, when published in the original studies, in Supplementary Figure 1.

#### Global meta-analysis

We analysed a total of 42,900 publicly available metagenomic samples (collectively >220Tbp) from 253 different studies, including the novel cohort of 447 samples from the citizen science project MyMicrobes described above (see **Supplementary Table 2**). The collection includes samples from 86 countries, 35 animal species and 6 different human body sites: gastrointestinal tract (stools, rectal swabs and biopsies), vagina, skin, oral cavity, respiratory tract and human milk. The selected environmental samples come from potentially faecally contaminated habitats, such as wastewater, freshwater, indoor surfaces, soil and polluted harbor marine waters.

### Public data download

Metagenomes publicly available on the 22^nd^ of April 2021 were downloaded using fetch-data (Coelho et al. 2021). The collection process aimed at leveraging the largest possible set of samples and to maximise the number of covered environments, with the only inclusion criteria being samples sequenced with whole genome sequencing on Illumina platforms. No minimum number of samples per study or minimum sequencing depth threshold was applied at this stage.

### Metadata curation

Manually curated metadata for each dataset include: health status, age, geography (country and continent name), westernised lifestyle or not, diagnosis for *C. difficile* infection (CDI), use of antibiotics, delivery mode, premature birth (pre-term or full-term), and sex. Subjects were categorised by age group, defined as follows: 0 ≤ infant ≤ 12 months, 1 year < child ≤ 10 years, 10 years < adolescent ≤ 18 years, 18 years < adult ≤ 65 years, and elderly > 65 years of age. If a specific age was not available, a range of age was provided in alternative. Given the broad range of perturbations in the gut microbiome associated with antibiotics intake, we ad-hoc defined “diseased” samples as any sample taken from a subject with any medically diagnosed disease or syndrome and/or intake of one or more antibiotics at the time of the sampling.

### Filtering on read count

Two consecutive read count filtering steps were performed on all samples: (i) samples with zero reads mapping to mOTU marker genes (Milanese et al. 2019) were discarded (n=1,091) and (ii) samples with less than 59 reads mapping to mOTU marker genes, corresponding to 95%ile calculated on the remaining samples, were discarded (n=2,075). An additional third read count filtering was performed only on human gut stool samples, corresponding to 99%ile, removing samples with less than 100 reads (n=132 samples). The resulting filtered dataset included 39,338 samples, of which 26,610 were human fecal metagenomes.

### Identification of timeseries-representative samples

Out of 26,610 samples, 24,690 had subject-level metadata and were associated with 12,352 unique subjects. For 3,573 subjects, multiple timepoints were available. The mean number of timepoints per subject was 2 (3.2 for infants, 1.7 for children, 3.5 for adolescents, 1.7 for adults and 1.3 for elderly), with a maximum of 205 time points per subject.

In order to avoid under- or over-estimating *C. difficile* prevalence, only one sample per time series was used in cross-sectional analyses. We distinguished three cases for each time series:

i. *C. difficile* observed in all samples. In this case, the sample with the highest *C. difficile* read count is selected as representative and the corresponding subject was considered *C. difficile* positive.
ii. *C. difficile* observed in none of the samples. In this case, the sample of the first time point was selected as representative and the corresponding subject was considered *C. difficile* negative.
iii. *C. difficile* observed in only some samples. In this case, the sample with the highest *C. difficile* read count was selected as representative and the corresponding subject was considered *C. difficile* positive.

1,920 samples had no subject identifier. In this case, we assumed one sample per subject. One representative sample for each time series was used for all downstream analyses, if not specified otherwise. The timeseries dereplication procedure described above has also been applied to the CDI study populations before downstream analysis.

### Metagenomic data processing

#### Taxonomic profiling

Metagenomes were taxonomically profiled at the species level with mOTUs v2.0 (Milanese et al. 2019), requiring the confident detection of at least two taxonomic marker genes. All data analyses were conducted in the R Statistical Computing framework v3.5 or higher. Only *C. difficile* positive samples were considered for *C. difficile* relative abundance estimation.

#### Microbiome diversity

Local community diversities (‘alpha’ diversities) of human gut samples were estimated by iteratively rarefying taxonomic count tables to 100 marker gene-mapping reads and computing average Hill diversities at q=0 (species richness), q=1 (exponential Shannon entropy) and q=2 inverse Simpson index), as well as evenness measures as ratios thereof. Unless otherwise stated, results in the main text refer to taxa richness. Differences in alpha diversity were tested using ANOVA followed by post hoc tests and Benjamini-Hochberg correction, as specified in the main text.

#### Prevalence and abundance estimations of C. difficile

Prevalence estimations of *C. difficile* over life time were based on human gut samples (stools, rectal swabs and biopsies) where the precise subject age in months was available (samples with missing or too broad or age ranges were discarded). Both *C. difficile* mOTUs2.0 (“ref_mOTU_0051” and “ref_mOTU_0052”) were considered in downstream analysis. Abundance estimations included only *C. difficile* human stool samples with precise age metadata.

### Species co-occurrence analysis

Human gut stool samples, with at least 100 reads per sample and with known age group and health status were considered for this analysis. One representative sample per time series was considered. Fisher’s exact test, followed by Benjamini-Hochberg correction, were applied to identify co-occurring species. Meaningful positive association was identified for species with adjusted p-value <0.05 and logarithm of the odds ratio >1, while meaningful negative association was identified for species with adjusted p-value <0.05 and logarithm of the odds ratio < -1.

### Machine learning modelling

L1-regularised LASSO logistic regression models to predict CDI status were built using the SIAMCAT R package (Wirbel et al. 2021) with 10-fold cross-validation. For this analysis, we focused on the subset of 10 CDI or diarrhoea-associated datasets (**Supplementary Table 1**) and then trained two different sets of models: one set of models to distinguish CDI samples and samples from healthy controls (excluding controls from diseased subjects) and another set of models to distinguish CDI samples and any type of control samples. In order to minimise overfitting and to counter batch effects (Wirbel et al. 2021), we pooled datasets across studies in a leave-one-study-out approach. In short, all except one study were jointly processed to train a LASSO model that was then used to predict the left-out study. Additionally, to check if the microbial signature for CDI was independent of *C. difficile*, we trained another set of models with the same cross-validation splits but excluded *C. difficile* from the feature table. Feature weights were extracted from the models, normalised by the absolute sum of feature weights, and averaged across cross-validation folds. For the heatmap in Figure 2, all microbial species that were assigned non-zero weights in at least 80% of cross-validation folds were included.

### Linear mixed-effect model

To test for differential abundance of microbial species between CDI and non-CDI samples while taking into account possible confounding factors, we employed linear mixed-effect models as implemented in the lmerTest package (Kuznetsova et al. 2017). After filtering for prevalence (prevalence of at least 5% in three or more studies), we tested the log-transformed abundance of each microbial species using a linear mixed-ffect model with “CDI status” as fixed and “Study” and “Age group” as random effects. Effect size and p-values were extracted from the model and p-values were corrected for multiple hypothesis testing using the Benjamini-Hochberg procedure. In addition, we used a linear mixed-effect model to test for a relationship between sequencing depth and the ability to detect *C. difficile* in CDI study populations as well as on the wider global set of samples (considering only dereplicated samples from time series). In this analysis, study was considered as a random effect.

### Pathogenic toxin genes detection in metagenomes

Identification of toxin-coding genes and their assignment was performed using the VFDB database (Liu et al. 2019), downloaded in March 2021. In our analysis we did not investigate the presence of the *C. difficile* binary toxin gene, since this toxin alone has not been associated with disease severity (Goldenberg and French 2011). Mapping against VFDB was performed via BWA (Li and Durbin 2009) and filtering via NGLess (Coelho et al. 2019), using 99% as minimal alignment percentage identity threshold and 75bp as minimal read length match. Toxin gene profiles were computed with gffquant (https://github.com/grp-bork/gff_quantifier, version 1.2.3) using the 1overN count model. Toxin genes included in VFDB refer to the following strains: *C. difficile* 630, *E. coli* CFT073, O44:H18 042 and O157:H7 str. EDL933, *S. flexneri* 2a str.301, *P. aeruginosa* PAO1, *C. perfringens* str.13 and SM101, *S. aureus* RN4220, subsp. aureus MW2 and N315. The comprehensive list of virulence-related genes for each of these strains is available on the VFDB website (http://www.mgc.ac.cn/cgi-bin/VFs/v5/main.cgi).

### Subspecies analysis

*C. difficile* subspecies detection was performed on all *C. difficile* positive samples with MetaSNV2 (Van Rossum et al. 2021). Reads from 2,428 samples were mapped against proGenomes v1 (Mende et al. 2017) species representatives genomes for 3 species in the Clostridium genus: i) specI_v2_0051 (NCBI taxonomy ID 272563, PRJNA78) *Clostridioides difficile*, ii) specI_v2_0052 (NCBI taxonomy ID 1151292, PRJNA85757) *Clostridioides difficile* and iii) specI_v2_1125 (NCBI taxonomy ID 1408823, PRJNA223331) *Clostridioides mangenotii.* By mapping to multiple genomes and only using the uniquely mapped reads, we focused on the species-specific core genomic regions. Mappings that had at least 97% identity and a match length of at least 45bp were kept. Mapping and filtering was performed using BWA (Li and Durbin 2009) and NGLess (Coelho et al. 2019). Reads that mapped uniquely across the 3 reference genomes were used to call single nucleotide variants (SNVs) using metaSNV2 with default parameters. Of the initial 2383 samples, 197 passed the requirements for robust SNV calling (8%) and 179 could be used to detect subspecies presence. Substructure within the population was assessed in the resultant SNV profiles according to a previously reported approach (Li and Durbin 2009; Costea et al. 2017). Briefly, dissimilarities between samples were calculated based on SNV abundance profiles and the resultant distance matrix was tested for clusters using the Prediction Strength algorithm (Yuan et al. 2020). Distance matrix is plotted using R and the pheatmap package with average clustering. Six samples with extreme dissimilarity to all other samples were removed from the distance matrix for illustrative purposes (SAMN08918181, SAMN09980608, SAMN10722477, SAMN13091313, SAMN13091317, SAMN13091322).

### Building SNP-tree

SNPs obtained via metaSNV2 were arranged into an alignment PHY input format file suitable for IQ-TREE2 tree-builder (Minh et al. 2020). Sites which had multiple variants/alleles (in this case only SNPs) were replaced with alternative non-reference allele if they were supported by >50% of reads mapped against specI_v2_0051 (NCBI taxonomy ID 272563, PRJNA78), see ‘Subspecies analysis**’** section for mapping details. This results in forced fixing of alleles across polymorphic sites within each sample. To include specI_v2_0052 (NCBI taxonomy ID 1151292, PRJNA85757) *Clostridioides difficile* and specI_v2_1125 (NCBI taxonomy ID 1408823, PRJNA223331) *Clostridioides mangenotii* into SNP input file for IQTREE2 we used Mauve(Darling et al., 2004) to 1) order contigs of both genomes against specI_v2_0051 genome, 2) align ordered contigs, 3) export SNP profile from Mauve and further merged alignment SNP profile on metaSNV2 output file using matching positions. Missing coverage was kept as gaps in the alignment and tree construction. Following command was used to build phylogenetic tree: iqtree -s >INPUT snp file< -m GTR+ASC -B 1000.

### Availability of data and materials

This study did not generate new unique reagents. Further information and requests for resources and reagents should be directed to and will be fulfilled by the lead contacts, Sebastian Schmidt and Peer Bork. Novel raw metagenomic sequencing data have been uploaded to the European Nucleotide Archive under the accession number PRJEB50977. The accession numbers relative to the other public studies included in this meta-analysis are available in Supplementary Table 2. The filtered taxonomic profiles and associated metadata used for the analyses are available in Supplementary Tables 4-6. The analysis code and commands used in the project are deposited in the GitHub repository at https://github.com/grp-bork/cdifficile_Ferretti_2022.

## Acknowledgements

We are thankful to the members of the groups of Peer Bork, Georg Zeller, Thomas Dandekar (University of Würzburg) and Dr. Athanasios Typas (EMBL) for fruitful discussions. We are also grateful to Lia Oken for her continuous support and her availability for brainstorming sessions. We acknowledge funding from EMBL and the European Research Council (MicrobioS grant no. ERC-AdG-669830 to P.B.). This work comprises results from Pamela Ferretti’s doctoral thesis.

## Author contributions

Conceptualization, P.F., T.SB.S. and P.B.; data collection, P.F., T.SB.S., R.A., A. F., W.A. and M.K.; metadata curation, P.F., T.SB.S., R.A., A.S., R.T., L.T, I.L. and M.K.; novel samples collection and processing, S. K., R.H. and A.T.; data analysis, P.F., J.W., O.M.M., T.VR., R.A., A. F., W.A., C.S. and T.SB.S.; writing – original draft, P.F. and T.SB.S.; writing – review & editing, P.F., J.W., O.M.M., T.VR, C.S., M.K, G.Z. T.SB.S. and P.B.; supervision, T.SB.S., G.Z. and P.B.; funding acquisition, P.B.

## Supplementary Figure Legends

**Supplementary figure 1.** (A) Prevalence of *C. difficile* and other antibiotic-associated diarrhoea (AAD) species, divided by study. For *C. difficile* only, the portion of samples with toxigenic *C. difficile* is shown (dotted segments). CDI diagnosis procedure is shown on top of each study. Each row represents the diagnostic algorithm (combination of tests) used to diagnose CDI. Tests performed are indicated with black dots (white if not). For example, in “Langdon et al. 2021”, CDI was diagnosed if symptoms were present and toxigenic culture for *C. difficile* was positive, or if symptoms were present and the enzymatic immunoassays for *C. difficile* was positive, or if pseudomembranous colitis was identified. Diagnostic protocols abbreviations: “EIA”: Enzymatic ImmunoAssays, “GDH”: glutamate dehydrogenase, “NAAT” nucleic acid amplification test (including PCR), “PMC”: pseudomembranous colitis. (B) Prevalence of the number of AAD species (*C. difficile* not included) identified in each CDI sample, divided by *C. difficile* positivity.

**Supplementary Figure 2**. Species-level composition for CDI, diseased control and healthy control samples as seen by mOTUs v2.0. Species with minimum relative abundance ≥ 0.01 and prevalence ≥ 0.1 are shown.

**Supplementary figure 3.** Species significantly enriched (in yellow) or depleted (in blue) in terms of relative abundance in CDI compared to diseased and healthy controls, as identified by linear mixed-effect model analysis. Species known to cause antibiotic-associated diarrhoea (AAD) are highlighted in red. *C. difficile* is not included in the analysis. Study and age group were considered as nuisance variables (and modelled as random effects). To be noted that *Enterobacteriaceae* sp. includes *Shigella flexneri* and *Escherichia coli*.

**Supplementary figure 4.** (A) To check the influence of sequencing depth on the detection of *C. difficile*, the marker gene coverage values across all samples were binned into ten quantiles. The proportion of *C. difficile*-positive samples is shown in the barplot. A possible association between coverage quantile and *C. difficile* detection was tested using a linear mixed-effect model with Study as random effect (P-value=0.63). (B) Adjusted p-values for ANOVA analysis aiming to explain variance in *C. difficile* presence by the available metadata, either as single factors (two top rows) or by a combination of multiple factors (sequential, bottom rows, in the exact order shown along the X axis), in data sets that were broken down by age group (infants versus adults).

**Supplementary figure 5.** (A) Geographical distribution across continents and countries for *C. difficile* prevalence in the stools of healthy infants (left) and adults (right). Only countries with *C. difficile* prevalence above 1% are shown. Prevalence was calculated based on presence/absence of *C. difficile* from mOTUs v2.0^30^ taxonomic profiles. (B) Prevalence of *C. difficile* across subjects diagnosed with specific diseases (left) and subjects taking antibiotics (right). Average *C. difficile* prevalence is shown with grey bars, age-group specific prevalence is highlighted with colored dots. Number of datasets used for prevalence estimations is reported in the lower part. Abbreviations: NEC: “Necrotizing enterocolitis”; ASCVD: “atherosclerotic cardiovascular disease”; T2D: “Type 2 diabetes”; CRC: “Colorectal cancer”; ADA: “advanced adenoma”; NAA: “non-advanced adenoma”. See Methods for details on age group definitions.

**Supplementary figure 6.** *C. difficile* prevalence in infants and children, divided by possible combinations of health status, prematurity and delivery mode.

**Supplementary figure 7.** Time of the first appearance of *C. difficile* in infants and children timeseries. The grey line indicates the total number of samples per age interval, considering all samples independently of health status, delivery mode, gestational age and diet.

**Supplementary figure 8.** Community similarity (Bray-Curtis index) of healthy infant-mother pairs in presence or absence of *C. difficile* in stools across the first four years of life. Only full-term infant samples were included. Mean comparison p-values calculated using t-test, * for P ≤ 0.05, ** for P ≤ 0.01, *** for P ≤ 0.001, **** for P ≤ 0.0001, empty when non significant.

**Supplementary figure 9.** Prevalence of *C. difficile* toxin genes in the stools of healthy and diseased subjects over lifetime, based on the detection of either one or both of *C. difficile* Toxin A (*tcdA*) or Toxin B genes (*tcdB*) (see Methods for details). Prevalence is calculated on the number of *C. difficile* positive samples. *C. difficile* toxin genes were exclusively found among human stool samples. See Methods for details on age group definitions.

**Supplementary figure 10.** Alpha diversity calculations in *C. difficile* positive and negative samples. (A) Species richness in healthy infants with detailed age resolution. (B-C) species richness in infant samples divided by delivery mode, gestational age and health status. (D) Species evenness (as the inverse Simpson index divided by Richness) across age groups and health status. Species richness for continents with at least five *C. difficile* positive samples are shown. The number of samples per group is shown under each boxplot. Mean comparison p-values calculated using t-test, * for P ≤ 0.05, ** for P ≤ 0.01, *** for P ≤ 0.001, **** for P ≤ 0.0001, empty when non significant. See Methods for details on age group definitions.

**Supplementary figure 11.** Species with co-occurrence (logOR>0) or co-exclusion (logOR<0) pattern with *C. difficile* across gut stool samples from humans (A) and animals (B). Labelled species are highlighted in aquamarine and violet, for humans and animals respectively. *Clostridium paraputrificum* and *Clostridium neonatale* were among the very few species significantly associated with *C. difficile* in both human and animal stools.

**Supplementary Figure 12.** Spearman’s Rho values for the species listed in Figure 5, across the two groups. Mean comparison p-values calculated using t-test, * for P ≤ 0.05, ** for P ≤ 0.01, *** for P ≤ 0.001, **** for P ≤ 0.0001, empty when non significant.

**Supplementary Figure 13.** SNV similarity across *C. difficile* positive samples from our global metagenomic survey. Tree rooted with *Clostridioides mangenotii* as outgroup. Missing metadata are shown in white. All metagenomes have ANI similarity >95%. See Methods for detailed analysis description.

## Supplemental Table Legends

**Supplementary Table 1.** Overview of the public metagenomic CDI study populations included in the CDI meta-analysis, divided by sample group.

**Supplementary Table 2.** Overview of the public metagenomic studies included in the global meta-analysis.

**Supplementary Table 3.** Mean *C. difficile* prevalence and relative abundance estimation across age groups and health status.

**Supplementary Tables 4.** Samples metadata before read filtering and timeseries dereplication steps.

**Supplementary Table 5.** Samples taxonomic profiles, as provided by mOTUs v2.0.

## Relevance of this study

To our knowledge, this is the first large-scale metagenomic meta-analysis on *Clostridioides difficile*. We investigated *C. difficile* prevalence in 42,900 metagenomic samples from 86 countries and used machine learning to identify a global CDI-specific microbial signature. Our findings suggest that *C. difficile* may be overdiagnosed in the clinical context, with other enteropathogens able to induce CDI-like symptomatology, erroneously contributing to the *C. difficile* burden estimation. Disentangling the real burden of *C. difficile* from other pathogens is a fundamental step towards the development of adapted therapeutic interventions. Furthermore, this study provides new insights into the gut microbiome composition in presence of *C. difficile,* not only in CDI patients, but also across other diseases and in healthy subjects. In particular, we found that *C. difficile* colonisation was associated with multiple indicators of a healthy gut microbiome development in infants, an age group in which *C. difficile* has been so far understudied. This study also represents a blueprint for future large-scale species-specific metagenomic meta-analyses.

